# A regimen compression strategy for commercial vaccines leveraging an injectable hydrogel depot technology for sustained vaccine exposure

**DOI:** 10.1101/2023.03.23.534005

**Authors:** Jerry Yan, Ben S. Ou, Olivia M. Saouaf, Emily L. Meany, Noah Eckman, Eric A. Appel

## Abstract

Equitable global access to vaccines requires we overcome challenges associated with complex immunization schedules and their associated economic burdens that hinder delivery in under resourced environments. The rabies vaccine, for example, requires multiple immunizations for effective protection and each dose is cost prohibitive, and therefore inaccessibility disproportionately impacts low- and middle-income countries. In this work we developed an injectable hydrogel depot technology for sustained delivery of commercial inactivated rabies virus vaccines. In a mouse model, we showed that a single immunization of a hydrogel-based rabies vaccine elicited comparable antibody titers to a standard prime-boost bolus regimen of a commercial rabies vaccine, despite these hydrogel vaccines comprising only half of the total dose delivered in the bolus control. Moreover, these hydrogel-based vaccines elicited similar antigen-specific T-cell responses and neutralizing antibody responses compared to the bolus vaccine. Notably, we demonstrated that while addition of a potent clinical TLR4 agonist adjuvant to the gels slightly improved binding antibody responses, inclusion of this adjuvant to the inactivated virion vaccine was detrimental to neutralizing responses. Taken together, these results suggest that these hydrogels can enable an effective regimen compression and dose-sparing strategy for improving global access to vaccines.

## 1. Introduction

The Global Vaccine Action Plan, enacted in 2011 by the World Health Assembly, made valuable efforts to promote more equitable access to existing vaccines.^[1–2]^ While great progress has been made, vaccine-preventable diseases still disproportionately affect some areas of the world. For example, polio has still not yet been globally eradicated despite polio vaccines first introduced in 1950s, and measles is undergoing an alarming resurgence. There are many underlying reasons for inequitable vaccination across the globe, which have been widely observed through the COVID-19 pandemic, including supply chain and distribution challenges.^[3]^ Moreover, many current vaccines have immunization schedules requiring multiple boosts, which can be extremely complicated based on supply and clinic access, both of which remain challenges for low- and middle-income countries (LMICs).

Rabies is one example of a disease that disproportionately affects rural communities in LMICs. Although commercial vaccines already exist for rabies (e.g., RabAvert, Imovax, etc.), they are not readily available to those in need. Rabies is responsible for roughly 60,000 human deaths annually around the world, and more than 95% of these deaths occur in Africa and Asia.^[4–5]^ Commercial rabies vaccines are inactivated viral vaccines, which consist of the whole virus with destroyed genetic material. Inactivated vaccines are unable to mimic natural infection due to their inability to replicate, therefore only persisting in the body for days rather than a period of weeks often observed in natural infection.^[6]^ As a result, the commercial rabies vaccines require multiple immunizations to mount an effective immune response, and regular serological testing is required to monitor protection, both of which are challenging to achieve in LMICs.

Developing technological approaches providing durable protection against severe disease with fewer required immunizations could be a promising avenue for improving vaccines for global health. One such approach is the sustained delivery of vaccine components, which has shown to improve humoral responses such as heightened antibody titers and improving neutralization potency. In a non-human primate model, Crotty et al. demonstrated that sustained delivery of an HIV envelope protein immunogen and a potent saponin-based adjuvant system from an implantable osmotic pump for over two weeks dramatically improved the magnitude and durability of binding antibody titer, and improved peak neutralization titers by over 20-fold, when compared to bolus administration of the same vaccine.^[7]^ The authors of this study demonstrated that sustained vaccine exposure prolonged germinal center reactions, which lasted for upwards of six months, indicating that this approach could potentially eliminate the need for booster immunizations in some indications. Unfortunately, the surgically implanted pumps used in this study are not clinically translatable for more infectious disease indications, highlighting a need for development of technologies capable of sustained delivery of vaccine components.

To address this challenge, several groups have developed microneedle technologies to simplify administration while achieving robust humoral and cellular immune responses, and in some cases even enabling sustained vaccine exposure.^[8–12]^ However, challenges associated with complex manufacturing and vaccine stability during typical manufacturing processes have hampered many of these technologies. Moreover, many of these technologies can’t make use of existing commercial vaccines. Our lab has previously developed a self-assembled hydrogel platform for biomedical applications including sustained delivery of physicochemically diverse therapeutic cargo.^[13–20]^ Our efforts have focused on the development of shear-thinning and self-healing physical hydrogels that can be easily injected and which can afford sustained delivery of multifarious vaccine cargo.^[13, 21–24]^ Specifically, these hydrogel materials leverage strong, non-covalent polymer-nanoparticle (PNP) interactions to crosslink biopolymer chains. The manufacturing of these materials is inexpensive and scalable, and we have previously shown that PNP hydrogel vaccines promote greater affinity maturation and more potent and durable humoral responses with several common subunit antigens for pathogens such as SARS-CoV-2 and influenza.^[13, 22–23]^ However, the versatility of the PNP hydrogel system has not been evaluated beyond subunit vaccines with, for example, inactivated virus vaccines such as the commercial vaccines for rabies.

In this work we sought to engineer the PNP hydrogel system to enable regimen compression for the commercial rabies vaccine RabAvert. We first characterized the hydrogels’ mechanical properties upon the addition of the inactivated virus vaccine cargo. We then evaluated humoral and cellular immune responses following sustained exposure from a single immunization of PNP hydrogel-based vaccines compared to a prime-boost series of bolus immunizations of the commercial vaccine. A single immunization of PNP hydrogel resulted in comparable antirabies virus IgG antibody titers and antigen-specific T cell responses. Moreover, a rabies virus neutralization test revealed that neutralizing responses from the single immunization PNP hydrogel-based vaccines were equivalent to those from the prime-boost bolus regimen. This study suggests that sustained delivery of clinical inactivated virus vaccines in PNP hydrogels induces durable neutralizing humoral immunity from a single immunization, potentially improving global access to these critical vaccines.

## 2. Results

### Hydrogel Development for Sustained Exposure of Commercial Vaccines

We have previously demonstrated that PNP hydrogels can act as a simple and effective platform for sustained delivery of subunit vaccines to improve humoral immune responses.^[13, 21–24]^ These hydrogels are formed by mixing hydrophobically-modified hydroxypropylmethylcellulose (HPMC-C_12_) and biodegradable nanoparticles made by nanoprecipitation of poly(ethylene glycol)-*b*-poly(lactic acid) block copolymers (PEG-PLA NPs). These materials are denoted PNP-X-Y, where X refers to the wt% loading of HPMC-C_12_ polymer and Y refers to the wt% loading of PEG-PLA NPs (*i.e*., PNP-1-10 comprises 1 wt% polymer and 10% NPs). Physical crosslinking in these hydrogels comprising entropy-driven dynamic interactions between the HPMC-C_12_ polymers and the surface of the PEG-PLA NPs as the polymers bridge between the NPs.^[25–27]^ This self-assembled, physically-crosslinked hydrogel system exhibits a high degree of shear-thinning, allowing for easy injection through standard needles and syringes, and rapid self-healing to form a robust subcutaneous depot.^[28–29]^ In this work, we engineered PNP hydrogels for controlled encapsulation and sustained delivery of commercial RabAvert rabies vaccine to achieve neutralizing immunity in a single immunization (**Figure 1a**).

**Figure 1.**
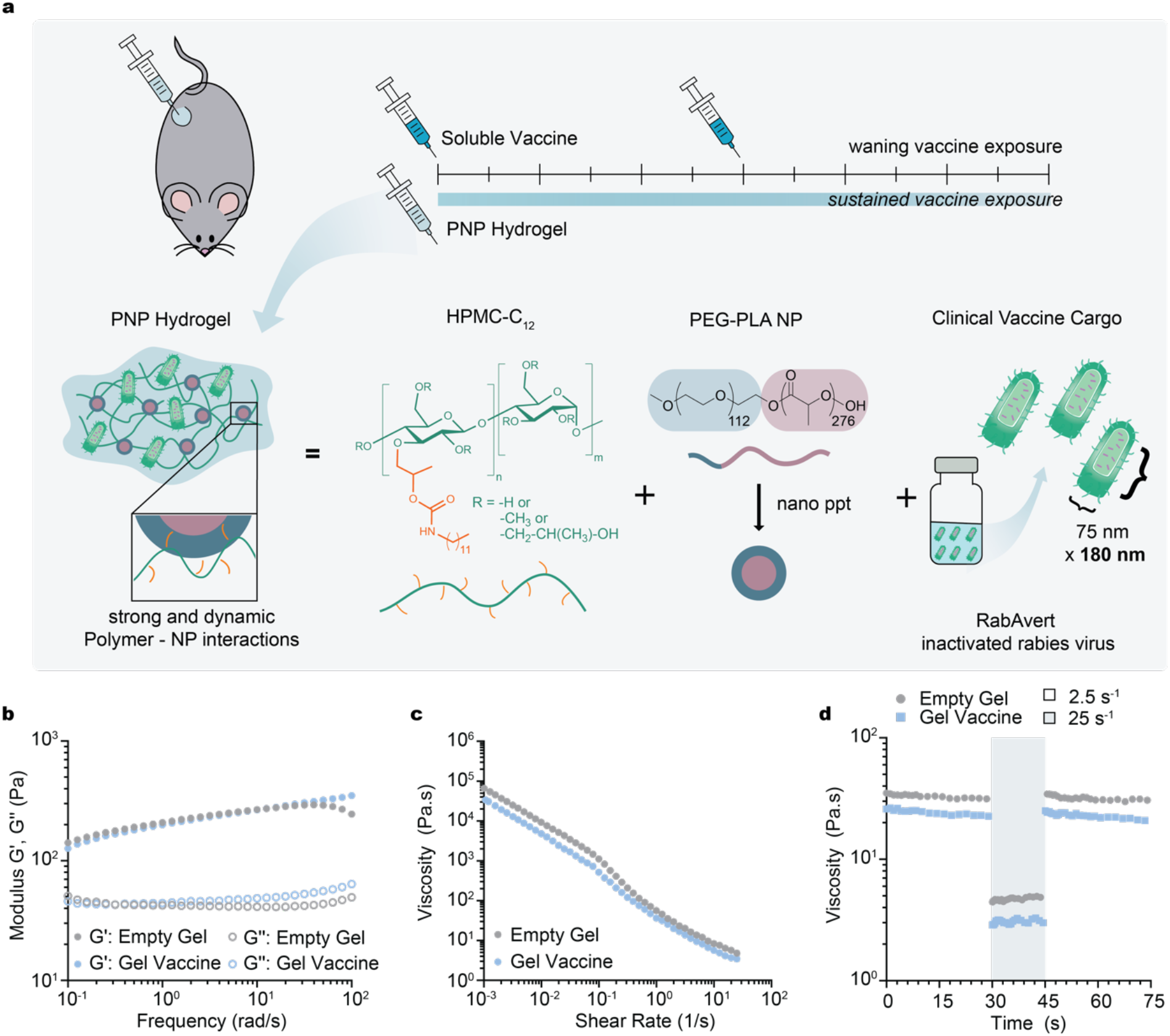
Sustained delivery of commercial vaccine cargo with an injectable depot technology enables a regimen compression and dose-sparing strategy eliciting durable immunity. **(a)**Schematic of injectable polymer-nanoparticle (PNP) hydrogel vaccines. Dodecyl-modified hydroxypropylmethylcellulose (HPMC-C_12_) is mixed with nanoparticles comprising poly(ethylene glycol)-*b*-poly(lactic acid) (PEG-PLA NPs) and commercially available RabAvert vaccine cargo to form vaccine-loaded PNP hydrogels. Dynamic, multivalent non-covalent interactions between the polymers and the NPs yield physically crosslinked hydrogels enabling sustained release of vaccine cargo. **(b)**Frequency-dependent oscillatory shear rheology and **(c)**shear-dependent flow rheology of empty and vaccine-loaded PNP hydrogel formulations indicate that vaccine incorporation doesn’t alter hydrogel properties. **(d)**Step-shear measurements of empty and vaccine-loaded PNP hydrogels over a cycle of alternating high shear (gray; 25 s^-1^) and low shear (white; 2.5 s^-1^) rates.

### Characterization of Hydrogel Shear-Thinning and Self-Healing Properties

We characterized the viscoelastic properties of PNP hydrogels encapsulating RabAvert to ensure that the inactivated rabies virus and other excipient components of the commercial vaccine did not interfere with rheological properties critical for injectability, self-healing, and depot formation. Frequency-dependent oscillatory shear experiments performed in the linear viscoelastic regime of the hydrogels showed that PNP-1-10 hydrogels comprising RabAvert had nearly identical frequency response compared to empty PNP-1-10 hydrogels (**Figure 1b**). For both formulations, the storage modulus (G’) remained above the loss modulus (G”) across the entire range of frequencies evaluated (i.e., tanδ < 1; **Figure S1**), indicating that these hydrogels exhibit solid-like properties necessary for robust depot formation.

To evaluate the injectability of these materials, a shear rate sweep was performed to characterize the hydrogels’ shear-thinning behaviors. The viscosity of the hydrogels decreased several orders of magnitude as the shear rate increased (**Figure 1c**). Step shear experiments were also performed to simulate the change in shear rate encountered during injection to evaluate the self-healing ability of these materials and their subsequent depot formation. Upon alternating between a high shear rate (25 s^-1^) and a lower shear rate (2.5 s^-1^), the viscosity of the hydrogel comprising RabAvert decreased by an order of magnitude under high shear, and rapidly (<5s) recovered to initial viscosity when the shear rate was decreased (**Figure 1d)**. These data demonstrate these hydrogel formulations can rapidly self-heal to form a robust depot following injection. Overall, these rheological behaviors confirmed that the inclusion of the commercial vaccine RabAvert does not interfere with the mechanical properties of the PNP hydrogels.

### Immune Responses to Vaccination

A standard dosage of RabAvert in humans is equivalent to at least 2.5 international units (IU) of potency.^[30]^ In published preclinical studies, doses varying from 1/5 to 1/20 of the human dose have been tested for protection in mouse models.^[31–34]^ To evaluate the humoral immune response to PNP hydrogel-based vaccines comprising RabAvert, we immunized C57BL/6 mice subcutaneously and collected sera over a twelve-week period (**Figure 2a**). Initial screening studies demonstrated three bolus immunizations of commercial RabAvert at days 0, 7, and 21 did not improve antibody titers compared to only two immunizations given at days 0 and 21 (**Figure S2**). These preliminary studies corroborate previous literature and recent updates to clinical vaccination recommendations that reduce the number of pre-exposure immunizations from three to two.^[35–39]^ Therefore, bolus commercial vaccine controls were administrated at day 0 (prime) and day 21 (boost) at a dose of 0.25 IU each for a total dose of 0.5 IU. In contrast, PNP-1-10 hydrogel-based vaccines were administrated at day 0 (prime only) at a total dose of 0.25 IU. In this way, hydrogel vaccine treatment groups received a single dose and while the control group received two doses of a standard bolus commercial vaccine. Additionally, a PNP-1-10 hydrogel vaccine group comprising monophosphoryl lipid A (MPLA) was evaluated to assess the effect of potent molecular adjuvants on the vaccine response of these hydrogel-based inactivated virus vaccines. MPLA is a potent TLR4 agonist used in commercial vaccine formulations for Hepatitis B, human papillomavirus, malaria, and shingles,^[40–41]^ and we have previously demonstrated that it provides excellent adjuvanticity in the context of PNP hydrogel-based subunit vaccines.^[13]^

**Figure 2.**
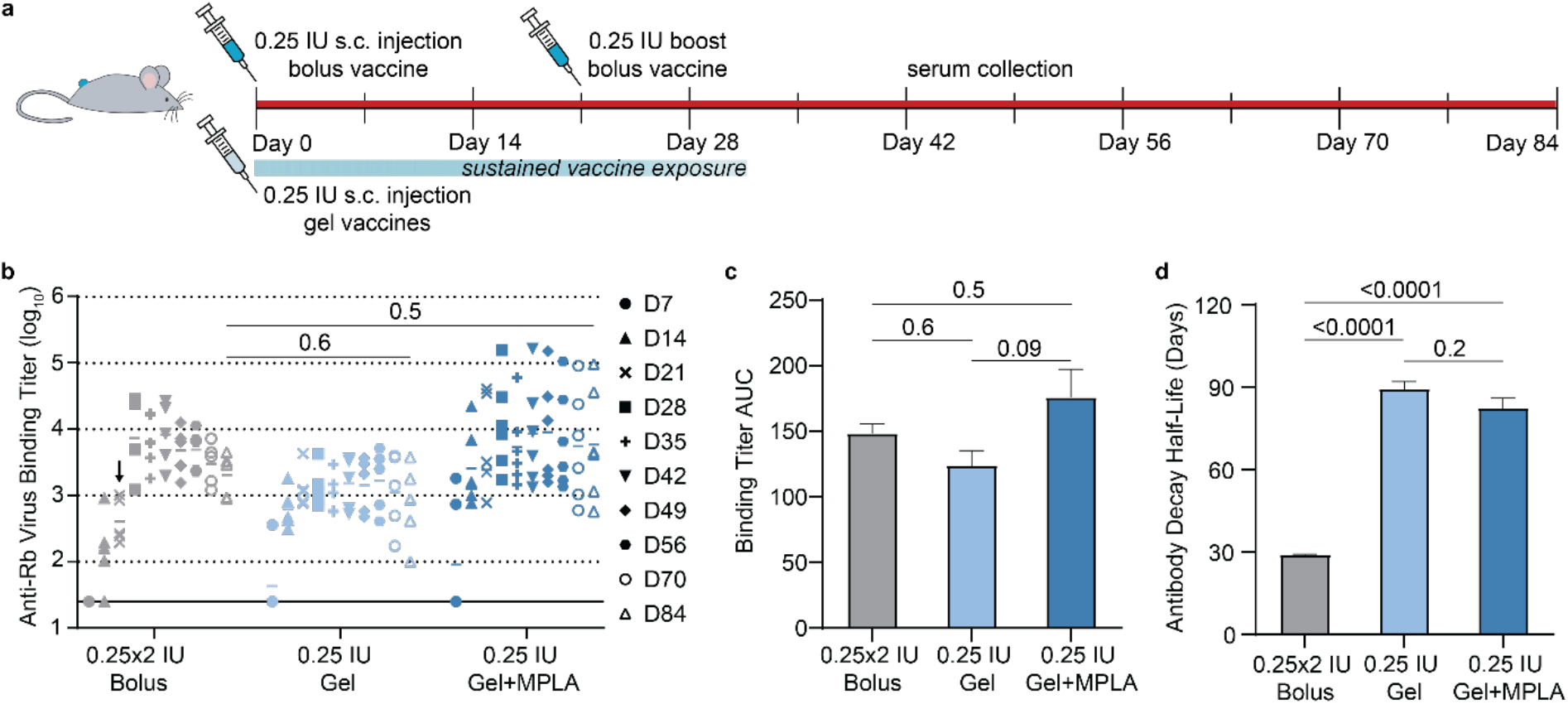
PNP-1-10 hydrogel vaccines induce durable humoral responses in mice with RabAvert. **(a)**Timeline of mouse immunizations and blood collection over a 12-week period. **(b)**ELISA antibody titers against the vaccine before and after boosting (arrow) of the bolus (n=5), the single immunization hydrogel (n=5), and the single immunization hydrogel comprising MPLA (n=6). Solid black line represents assay limit of detection and statistical comparisons were made with data from D84. **(c)**AUC data calculated from the longitudinal titers collected across the 12-week timeframe. **(d)**Antibody decay half-life calculated from D28 through D84 antibody titers using a parametric bootstrapping method to fit to an exponential decay model. Data are shown as mean +/- SEM. *p* values were determined using the linear model in JMP and Tukey’s HSD post-hoc test.

Following immunization, we quantified total IgG antibody titers against the RabAvert vaccine over the entire twelve-week period (**Figure 2b**). We observed that one injection of 0.25 IU PNP-1-10 hydrogel vaccines, both with and without MPLA, resulted in seroconversion across all treated animals by day 14. In contrast, not all mice in the bolus vaccine group had detectable antibody titers at this same timepoint. After the bolus control received a boost on day 21, all three groups were observed to maintain detectable titers over the entire twelve-week period. Notably, there was no significant difference in antibody titers between the two-dose bolus control receiving a total dose of 0.5 IU and single-dose of hydrogel vaccine groups receiving a total dose of 0.25 IU, observed at the latest timepoint of the study (day 84). To quantify total antibody exposure over the twelve-week period, we calculated the area under the curve (AUC) for all groups (**Figure 2c**). There were no significant differences observed in AUC values across the three treatment groups, demonstrating that a single hydrogel immunization with or without MPLA adjuvant elicited similar total antibody exposure across a twelve-week period compared to two immunizations of bolus vaccine. The single hydrogel immunizations thus demonstrated rapid seroconversion and durable potency, despite only containing half the doses of a traditional bolus vaccine.

We have reported in previous work that it is the formation of a persistent hydrogel depot that slowly dissolves away that provides control of sustained cargo release.^[28]^ In the current study, we tracked the hydrogel depot persistence time for the hydrogel vaccine groups. The hydrogel depot persisted for 23-28 days, averaging 26 days for half of the cohort of treated animals (**Figure S3**). This range of persistence time is notably longer than the total exposure period of the bolus vaccine, where the boost is administrated at day 21, and consistent with previous observations made with empty hydrogels.^[28]^ These observations suggest that hydrogel erosion and cargo release is dominated by hydrogel formulation selection and irrespective of the cargo.

To evaluate the durability of antibody responses, we fit an exponential decay model via bootstrapping to the antibody titer data after the peak observed a day 28. From this model, we estimated the half-life of decay for binding antibody titers for each treatment group (**Figure 2d**). The estimated half-life for the prime-boost bolus control group was found to be 28.6±0.2 days, while the half-lives observed for hydrogel-based vaccines with and without MPLA were found to be significantly longer at 82±4 days and 89±4 days, respectively (mean±s.e.m.; *p* < 0.0001 for either gel treatment compared with the bolus control). These data demonstrate that the hydrogel depot technology elicits dramatically prolonged antibody titers following a single administration, suggesting these hydrogel-based vaccines enable regimen compression and dose-sparing that can potentially improve global access.

In addition to the magnitude and durability of total IgG titers, we evaluated the IgG isotypes by ELISA to determine if the hydrogel vehicle altered the effects of RabAvert vaccine on immune signaling. Specifically, the ratio of IgG2c to IgG1 titers is a useful metric for Th1 versus Th2 skewing.^[42]^ We assessed titers for the bolus and gel treatment groups at week six, three weeks after the boost given in the bolus control group (**Figure 3a-c**). Notably, one of the mice in the bolus treatment group did not have detectable IgG1 antibody titer. On the other hand, all the mice receiving a single immunization hydrogel-based vaccine achieved seroconversion for IgG1 antibodies. The bolus control commercial vaccines exhibited an IgG2c/IgG1 ratio below 1, suggesting skewing slightly towards a Th2 response (**Figure 3d**). The hydrogel-based vaccine group elicited a comparable response, exhibiting a comparable IgG2c/IgG1 ratio and maintaining the skewing towards a Th2 response, consistent with previous observations from our lab.^[13]^ The hydrogel-based vaccine comprising MPLA exhibited the highest IgG2c/IgG1 ratio, corroborating previous literature suggesting that potent TLR agonists such as MPLA skew towards a cellular-mediated Th1 response.^[43]^

**Figure 3.**
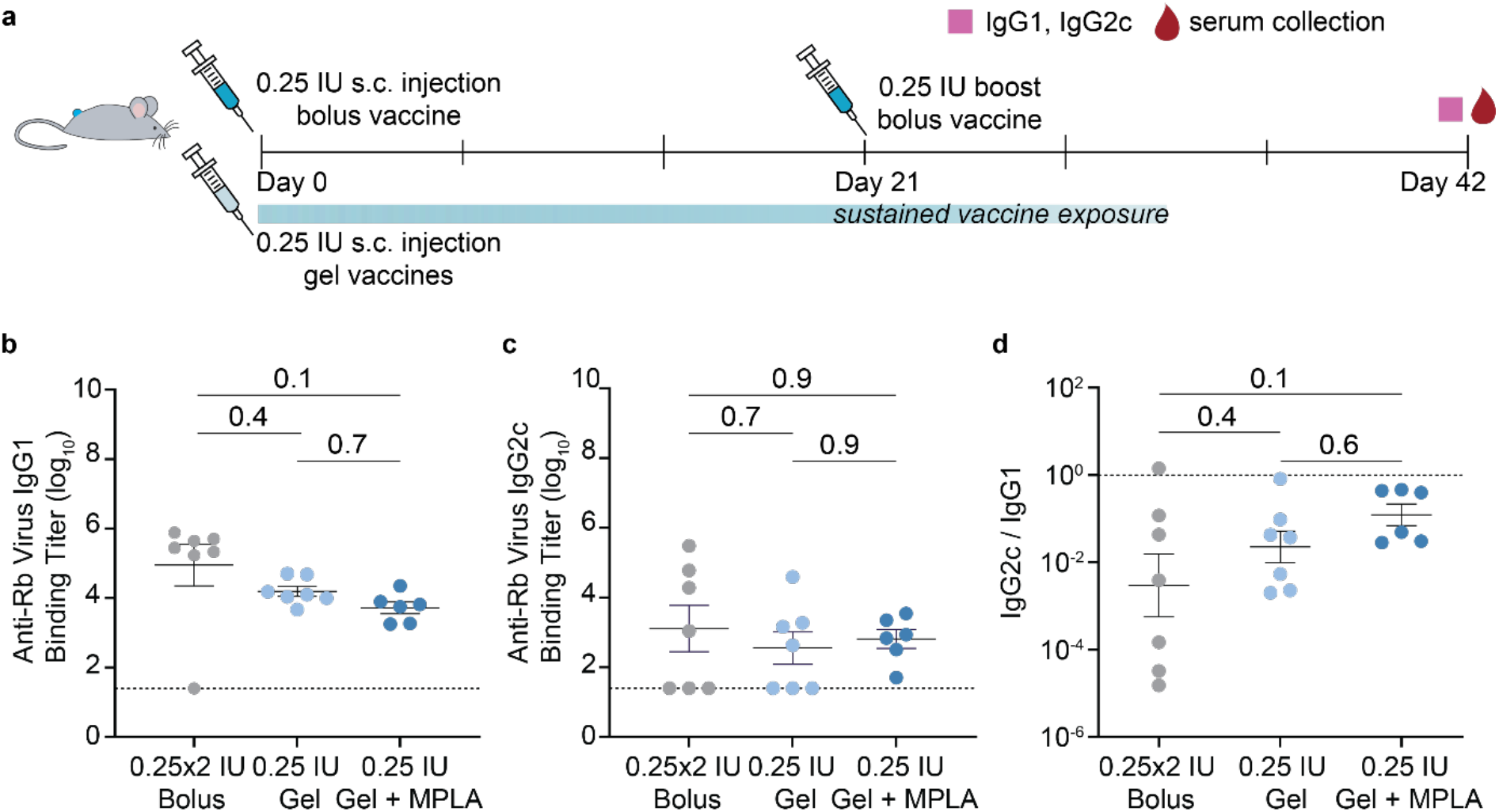
Single immunization hydrogel vaccine elicits comparable IgG2c and IgG1 antibody titers and overall skewing. **(a)**Timeline of mouse immunizations and blood collection at week 6 to determine IgG subtype titers for the bolus (n = 7), hydrogel vaccine (n = 7), and hydrogel vaccine with MPLA (n = 6). Serum anti-inactivated Rb virus **(b)**IgG1 titers, **(c)**IgG2c titers and **(d)**ratio of IgG2c to IgG1 titers. Lower ratios suggest a Th2-skewed humoral response while higher values suggest a Th1-skewed cellular response. Data are shown as mean +/- SEM. *p* values were determined using the linear model in JMP and Tukey’s HSD post-hoc test.

### Characterization of Cellular Infiltration into the Hydrogel Niche

Based on the differences observed in IgG2c/IgG1 ratio for hydrogel-based vaccines with and without MPLA, we sought to understand the potential impact of the adjuvant on the hydrogel the creation of a local inflammatory niche within the hydrogels. We have previously demonstrated that PNP-2-10 hydrogels can both retain subunit vaccine cargo over prolonged timeframes and enable infiltration of endogenous immune cells.^[21, 24]^ To better understand this hydrogel niche and its effect on immune responses, we evaluated the degree and composition of cellular infiltrate into the hydrogel microenvironment for both vaccine-loaded hydrogel groups (**Figure 4**; **Figure S4**). Seven days following subcutaneous injection, hydrogel depots were excised from the subcutaneous tissue and the cellular infiltrate was evaluated with flow cytometry (**Figure 4a**). Both hydrogel treatment groups experienced a large influx of immune cells, although the addition of MPLA to the hydrogel-based vaccines resulted in almost 2-fold higher recruitment of CD45^+^ leukocytes and over 3-fold higher recruitment of CD11b^+^CD11c^-^ myeloid cells. Overall, diverse cell populations were observed in vaccine-loaded hydrogels, both with and without additional MPLA adjuvant, including dendritic cells, B-cells, monocytes, and other myeloid and non-myeloid cells (**Figure 4b-g**).

**Figure 4.**
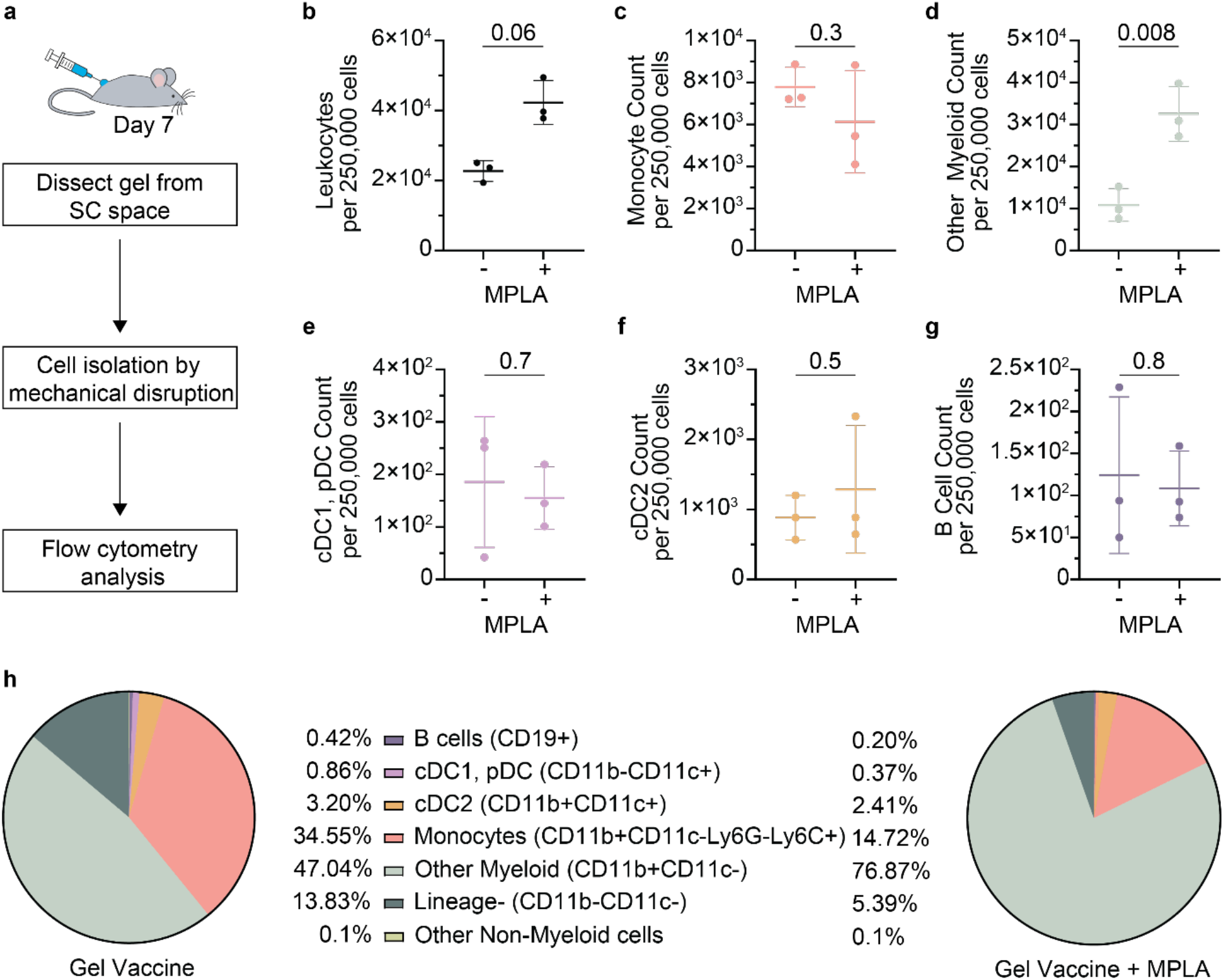
Single administration hydrogel vaccines adjuvanted with MPLA attract a diverse set of cell types to the hydrogel niche. **(a)**Schematic of experimental workflow: hydrogels (n = 3 each) were excised seven days after injection, mechanically dissociated, and analyzed via flow cytometry. **(b)**Average percentage of major immune subsets among total CD45^+^ infiltrate, with or without MPLA adjuvant. **(c-j)**Flow cytometry counts of live cells, CD45^+^ leukocytes, and CD45^+^ immune subsets. *p* values are determined by a Student’s T-test. Table S1 lists all fluorophores used for gating and analysis.

While the addition of MPLA significantly increased the number of myeloid cells recruited to the hydrogel niche, the total number of monocytes recruited did not significantly change with the addition of MPLA. As a result, monocytes composed a significantly lower proportion of the cellular response for the MPLA-adjuvanted hydrogel niche compared to the hydrogel niche comprising only the inactivated rabies virus vaccine (**Figure 4h**). Moreover, while monocytes comprised one of the largest cellular subsets for both gel niches, monocytes made up 35% of the cellular infiltrate in the vaccine-loaded hydrogel niche but only 15% of the MPLA-adjuvanted vaccine-loaded hydrogel niche. Based on our previous work demonstrating antigen uptake by infiltrating monocytes,^[24]^ these cell populations could potentially be playing a role in antigen presentation and subsequent germinal center responses leading to more sustained humoral responses.^[44–45]^ Overall, while there was substantial cellular infiltration into both hydrogel systems, the addition of MPLA not only increased the number of leukocytes infiltrating, but also drastically altered the overall composition of infiltrating cells.

### Rabies Viral Neutralization Assay

After confirming that a single immunization of hydrogel-based vaccines generated comparable antibody titers to a prime-boost regimen of a bolus clinical vaccine, we sought to determine the neutralizing activity of the sera. The rapid fluorescent foci inhibition test (RFFIT) involves serum-mediated inhibition of viral entry into cultured cells.^[33, 46]^ We tested viral neutralizing antibody titers from sera of individual mice at day 42 by evaluating the endpoint titer from the range of sera concentrations for half-maximal inhibition of infectivity (**Figure 5a**). According to World Health Organization (WHO) guidelines, RFFIT endpoint titers above 0.5 IU/mL are strongly correlated with preventing disease from an otherwise lethal challenge of rabies virus.^[47]^ With this endpoint titer threshold for “protection”, a prime-boost regimen of the commercial RabAvert vaccine and a single administration regimen of hydrogel-based vaccine resulted in comparable protection, achieving full protection in 55% and 47% of immunized mice, respectively (*p* = 0.6; **Figure 5b**). Notably, the single administration hydrogel vaccine comprising MPLA adjuvant performed only provided full protection in 17% of immunized mice. A majority of the MPLA-adjuvanted hydrogel vaccine cohort were significantly below the endpoint titer threshold for protection (**Figure S5**). These observations demonstrated that a single administration of PNP-1-10 hydrogel-based vaccine provided comparable neutralizing responses to the standard prime-boost bolus vaccines. On the other hand, the inclusion of MPLA adjuvant in the hydrogel weakened the neutralizing activity of the sera.

**Figure 5.**
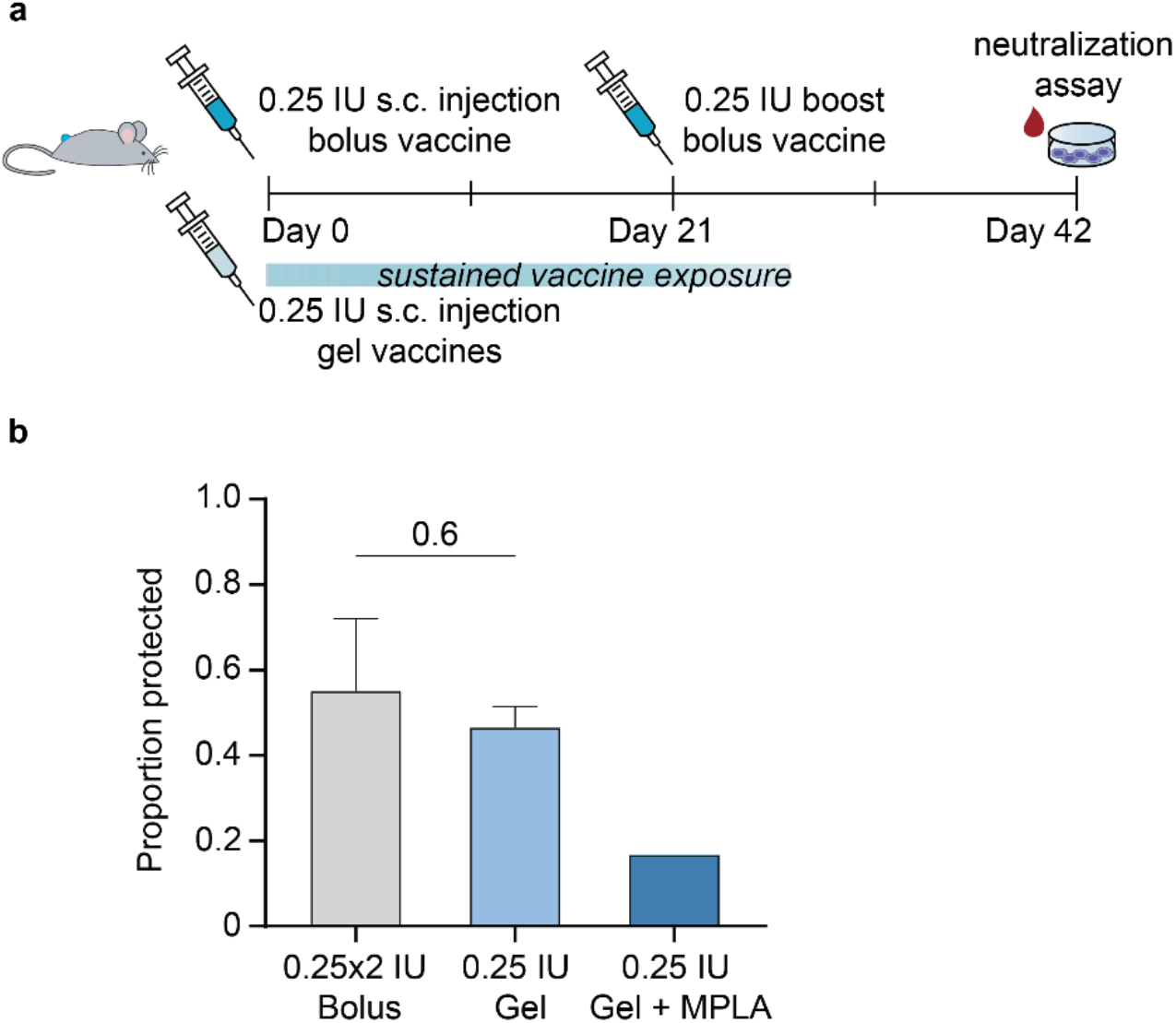
Single administration hydrogel vaccine without adjuvant elicited comparable neutralization titers to a standard prime-boost vaccine regimen. **(a)**Timeline of the experimental setup and terminal serum collection at week 6 to conduct neutralization assay RFFIT. **(b)**Proportion of mice fully protected in neutralization assay from two independent experiments for the bolus group (n = 6, n = 7), hydrogel group (n = 6, n = 7), and one independent experiment for the hydrogel with MPLA group (n = 6). Serum samples with over 0.5 IU/mL rabies virus neutralizing antibody titer are considered fully protected by WHO standards. *p* value was determined using a Student’s t-test.

### Cell-Mediated Immune Responses

Since a prime-boost bolus regimen of RabAvert is known to produce effective short-term protection from infection and severe disease partially mediated through cellular immunity,^[48]^ we sought to also evaluate the T cell responses elicited by hydrogel-based vaccines. We only evaluated responses of hydrogel-based vaccines without MPLA as the poor neutralization results indicated that this vaccine is irrelevant. To assess cellular responses, we isolated splenocytes at day 42 (three weeks after the bolus control group received the boost dose) and stimulated with rabies virus glycoprotein peptides for antigen-specific ELISpot (**Figure 6a**). As the most abundant protein on the viral surface, the rabies glycoprotein is a main target for vaccine-induced immunity.^[49–50]^ Following peptide stimulation, we measured IFNγ secreting CD8^+^ T cells and signal was observed across both vaccine groups (**Figure 6b**). A single administration of vaccine-loaded hydrogel induced a similar frequency of activated CD8^+^ T cells compared to two administrations of standard bolus vaccine (**Figure 6c**), indicating that hydrogel-based sustained vaccine delivery did not significantly alter T cell responses.

**Figure 6.**
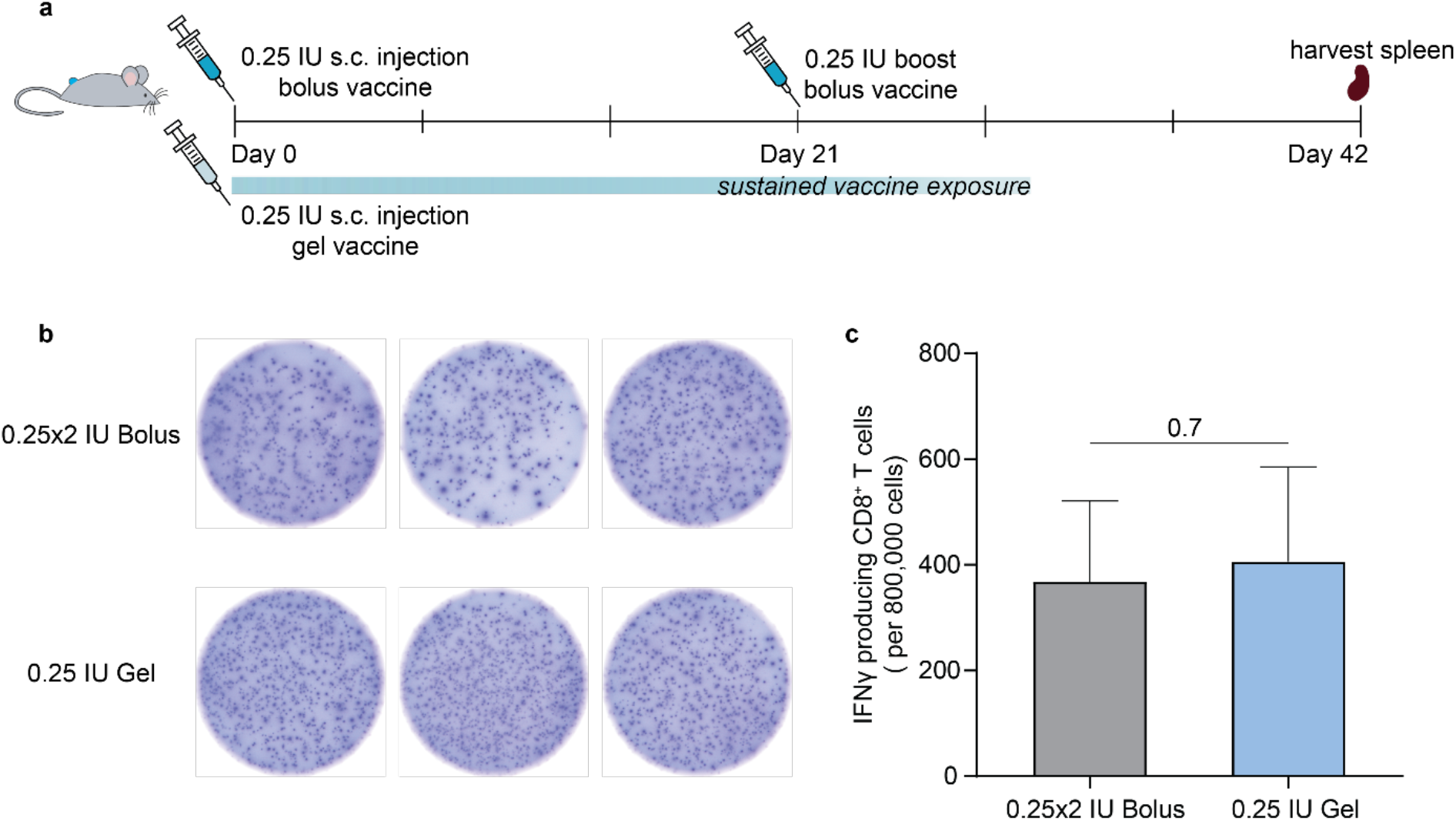
Single administration hydrogel vaccines do not alter antigen-specific CD8^+^ T cell responses. **(a)**Timeline of the experimental setup and spleen collection at week 6 to determine antigen-specific CD8^+^ T cell population. **(b)**Representative images of visible spots from IFNγ producing CD8^+^ T cell populations after simulation with 10 μg of rabies glycoprotein peptide. **(c)**The number of IFNγ producing CD8^+^ T cells upon antigen stimulation of 800,000 splenocytes/well for the bolus (n = 7) and hydrogel (n = 6) treatment groups. Data are shown as mean +/- SEM. *p* value was determined using a Student’s t-test.

## 3. Discussion

RabAvert, like other inactivated virus vaccines, requires multiple boosts to acquire sufficient neutralizing immunity. Such high burden, both in the burden of the immunization schedule and costs associated with vaccination, limits the accessibility of these vaccines in under resourced areas. In this work we sought to demonstrate that an injectable hydrogel depot technology capable of sustained vaccine exposure can form the basis for a single-administration immunization strategy capable of eliciting durable, protective immune responses. We have previously shown that PNP hydrogels can improve the potency and durability of humoral immune responses to subunit vaccines.^[13, 21–24]^ We hypothesized that this simple-to-implement self-assembled hydrogel technology could similarly improve the durability of immune responses through controlled delivery of commercial vaccines such as RabAvert, an inactivated virus vaccine. Unlike our previous work with subunit antigens, commercial vaccines like RabAvert include various excipients relevant for the manufacturing and stability of the drug product. The effect of these additional vaccine excipients on PNP hydrogel formation have not previously been studied. Here we found that the PNP hydrogels maintained similar viscoelastic rheological behaviors, shear-thinning behaviors, and injectability upon the addition of RabAvert. The simplicity of mixing commercial vaccines into the PNP hydrogels demonstrates the potential for their use as a platform strategy for improving diverse types of vaccines.

A single immunization of RabAvert-loaded hydrogel vaccine elicited comparable antibody titers to a prime-boost immunization series of bolus RabAvert. It is important to highlight that the total dose in the hydrogel was half of that delivered in the bolus control vaccines (0.25 IU compared to 0.5 IU), demonstrating the potential for dose-sparing. Furthermore, hydrogel-based vaccines demonstrated more durable antibody responses, with an estimated antibody decay half-life three-fold higher than the bolus vaccine control. Additionally, a single immunization of the hydrogel-based vaccines was observed to maintain the humoral immunological profile of the RabAvert vaccine, most notably in the Th2/Th1 antibody subtype skewing. We observed a slight skewing towards Th2 antibody-mediated responses for both bolus and hydrogel-based vaccines, consistent with prior literature.^[34, 51]^ Furthermore, we also observed no significant difference in the proportion of mice which could be characterized as “protected” with rabies virus neutralizing antibody titers above the WHO threshold in a well-characterized neutralization assay. Future work will uncover whether the humoral and cellular immune responses observed in this study over a 12-week period can be maintained for longer periods of time, perhaps by evaluating antibody titers and neutralization titers over longer timeframes, as well as characterizing various memory cell populations.

Prior to the present study, we had not evaluated cellular immunity induced by sustained vaccine delivery from our PNP hydrogel vaccines. Cellular-mediated responses have been positively correlated with levels of viral neutralizing antibody titers in rabies vaccination and providing long-term protection,^[52–54]^ though the exact contribution of CD8^+^ T cell response for protection against rabies remains unclear.^[51–52]^ Here we demonstrated that T cell responses were comparable between prime-boost bolus and single-administration hydrogel-based immunization regimens. Thus, while the PNP hydrogel system provide benefits in the form of more durable responses through sustained delivery of vaccine cargo, it does not appear to alter the flavor of immune responses imparted by the vaccine itself.

In addition to evaluating the ability of our PNP hydrogels to improve vaccine responses with a commercial vaccine, we also evaluated the impact of including a potent TLR4 agonist adjuvant, MPLA, currently used in several commercial vaccine products. A single administration of these MPLA-adjuvanted hydrogel-based vaccines elicited similarly sustained antibody titers over the twelve-week monitoring period, as well as a similar Th2/Th1 skewing. However, perhaps surprisingly, there was a drastically lower level of protection observed in neutralization assays. While the addition of MPLA elicted robust binding antibody titers, it dramatically lowered neutralizing antibody titers compared to hydrogel-based vaccines without MPLA added. We hypothesize that the impact MPLA has on altering the magnitude and composition of cellular infiltrate in the local hydrogel niche negatively affects downstream immune responses. We found that while MPLA inclusion resulted in more cells infiltrating the hydrogel niche, it also resulted in a significantly lower proportion of monocytes amongst infiltrating cells within the hydrogel niche. As monocytes have been previously implicated in antigen presentation and subsequent germinal center responses, leading to improved humoral immune responses,^[44–45]^their reduced presence in the local inflammatory niche may be responsible for weaker neutralizing responses.

It is important to highlight that the molecular mechanisms of many available adjuvants are not well understood.^[55]^ In this regard, other literature has shown that adjuvanted vaccines in mice can induce generation of long-lived plasma cells and neutralizing antibody titers, which may be crucial for long-term immunity.^[38, 55–56]^ Furthermore, several adjuvanted inactivated virus vaccines have entered phase 3 clinical trials and have shown promise in generating robust seroprotection.^[57–58]^ There is therefore, a rich landscape for further exploration of the impact of the addition of molecular adjuvants to hydrogel-based vaccines of the type we describe here on downstream immune responses.

Overall, approaches to overcome the challenge of equitable global accessibility must necessarily include technological development, either through novel modalities such as mRNA vaccines or depot technologies for sustained vaccine delivery. While sustained vaccine delivery technologies have mainly been studied in subunit vaccines for rapidly mutating viral pathogens such as HIV, influenza, and SARS-CoV-2,^[7, 59–60]^ we demonstrate that they show promise for immunization regimen compression and dose sparing with commercially available inactivated virus vaccines.

## 4. Conclusions

In summary, we report the development of a PNP-1-10 hydrogel platform enabling facile encapsulation and sustained delivery of a commercial inactivated virus vaccine, eliminating the need for boosts while maintaining a similar immunological profile. Compared to double the total dosage received in a standard prime-boost bolus vaccine regimen, single administration PNP hydrogel vaccines maintained similar antibody titers over three months, matched Th1/Th2 skewing, produced consistent cellular-mediated immunity, and generated comparable neutralization protection against rabies virus infection. This robust single immunization strategy has the potential to help overcome the challenges associated with global accessibility of commercial vaccines by both reducing the burden of multiple immunizations and reducing costs of vaccine delivery through dose sparing.

## 5. Methods

### Materials

HPMC (meets USP testing specifications), N,N-diisopropylethylamine (Hunig’s base), hexanes, diethyl ether, N-methyl-2-pyrrolidone (NMP), dichloromethane (DCM), lactide (LA), 1-dodecylisocynate, and diazobicylcoundecene (DBU) were purchased from Sigma-Aldrich and used as received. Monomethoxy-PEG (5 kDa) was purchased from Sigma-Aldrich and was purified by azeotropic distillation with toluene prior to use.

### Preparation of HPMC-C_12_

HPMC-C_12_ was prepared according to previously reported procedures.^22, 30, 52^ HPMC (1.0 g) was dissolved in NMP (40 mL) by stirring at 80°C for 1 h. Once the solution reached room temperature (RT), 1-dodecylisocynate (105 mg, 0.5 mmol) and N,N-diisopropylethylamine (catalyst, ~3 drops) were dissolved in NMP (5.0 mL). This solution was added dropwise to the reaction mixture, which was then stirred at RT for 16 h. This solution was then precipitated from acetone, decanted, redissolved in water (~2 wt %), and placed in a dialysis tube for dialysis for 3-4 days. The polymer was lyophilized and reconstituted to a 60 mg/mL solution with sterile PBS.

### Preparation of PEG-PLA NPs

PEG-PLA was prepared as previously reported.^[21, 61–62]^ Monomethoxy-PEG (5 kDa; 0.25 g, 4.1 mmol) and DBU (15 μL, 0.1 mmol; 1.4 mol % relative to LA) were dissolved in anhydrous dichloromethane (1.0 mL). LA (1.0 g, 6.9 mmol) was dissolved in anhydrous DCM (3.0 mL) with mild heating. The LA solution was added rapidly to the PEG/DBU solution and was allowed to stir for 10 min. The reaction mixture was quenched and precipitated by a 1:1 hexane and ethyl ether solution. The synthesized PEG-PLA was collected and dried under vacuum. Gel permeation chromatography (GPC) was used to verify that the molecular weight and dispersity of polymers meet our quality control (QC) parameters. NPs were prepared as previously reported.^[21, 61–62]^ A 1 mL solution of PEG-PLA in acetonitrile (50 mg/mL) was added dropwise to 10 mL of water at RT under a high stir rate (600 rpm). NPs were purified by centrifugation over a filter (molecular weight cutoff of 10 kDa; Millipore Amicon Ultra-15) followed by resuspension in PBS to a final concentration of 200 mg/mL. NPs were characterized by dynamic light scattering (DLS) to find the NP diameter, 37 ± 4 nm.

### PNP Hydrogel Preparation

The hydrogel formulation contained 1 wt% HPMC-C_12_ and 10 wt% PEG-PLA NPs in PBS. These gels were made by mixing a 1:3:2 weight ratio of 6 wt% HPMC-C_12_ polymer solution, 20 wt% NP solution, and PBS containing all other vaccine components. The NP and aqueous components were loaded into one syringe, the HPMC-C_12_ was loaded into a second syringe and components were mixed using an elbow connector. After mixing, the elbow was replaced with a 21-gauge needle for injection.

### Material Characterization

Rheological characterization was performed on PNP hydrogels using a TA Instruments Discovery HR-2 torque-controlled rheometer (TA Instruments) fitted with a Peltier stage. All measurements were performed using a serrated 20 mm plate geometry at 25°C with a 500 μm gap height. Dynamic oscillatory frequency sweep measurements were performed from 0.1 to 100 rad/s with a constant oscillation strain in the linear viscoelastic regime (1%). Steady shear experiments were performed by alternating between a low shear rate (1 s^-1^) and high shear rate (10 s^-1^) for 30 seconds and 15 seconds, respectively. Shear rate sweep experiments were performed from 25 s^-1^ to 0.001 s^-1^.

### Vaccine Formulations

The vaccines contained a 0.25 IU dose of RabAvert (GlaxoSmithKline) in 100 μL hydrogel or PBS based on the treatment group. For the bolus vaccines, the above vaccine doses were prepared in PBS and loaded into a syringe for administration. For the PNP hydrogels, the vaccine cargo was added at the appropriate concentration into the PBS component of the gel and combined with the NP solution before mixing with the HPMC-C_12_ polymer, as described above.

### Mice and Vaccination

C57BL/6 mice were purchased from Charles River and housed at Stanford University. 8-10 week-old female mice were used. Mice were shaved prior to initial immunization. Mice received 100 μL hydrogel or bolus vaccine on their backs under brief isoflurane anesthesia. Bolus treatments were injected with a 26-gauge needle and hydrogels were injected with a 21-gauge needle. Mouse blood was collected from the tail vein for survival bleeds over the course of the study.

### Mouse Serum ELISAs

Anti-rabies inactivated virus antibody titers were measured using an end-point ELISA. 96-well Maxisorp plates (Thermo Fisher) were coated with whole rabies inactivated virus (RabAvert, GlaxoSmithKline) at 0.5 IU/mL in 1X PBS (pH 7.4) overnight at 4°C. Plates were then blocked with 1% non-fat dry milk (Rockland) for 1h at RT. Serum samples were serially diluted starting at a 1:100 dilution in 1% bovine serum albumin (BSA in 1x PBS) and incubated on blocked plates for 2h at RT. One of the following goat-anti-mouse secondary antibodies was used: IgG Fc-HRP (1:10,000, Invitrogen A16084), IgG1 heavy chain HRP (1:10,000, abcam ab97240), or IgG2c heavy chain HRP (1:10,000, abcam ab97255). The secondary antibody was added at the dilution listed (in 1% BSA) for 1 h at RT. 5X PBS-T washes were done between each incubation step. Plates were developed with TMB substrate (TMB ELISA Substrate (High Sensitivity), Abcam). The reaction was stopped with 1 M HCl. Plates were analyzed using a Synergy H1 Microplate Reader (BioTek Instruments) at 450 nm. The total IgG and the subtypes were imported into GraphPad Prism 8.4.1 to determine the endpoint titers by fitting the curves with a five-parameter asymmetrical non-linear regression at a cutoff of 0.1. Samples failing to calculate endpoint threshold at a 1:100 dilution were set to a titer cutoff of 1:25 or below the limit quantitation for the assay. *p* values listed were determined using a Student’s t-test on GraphPad Prism software.

### Antibody Half-life Decay Model

To compare the titer half-lives, a parametric boostrapping method was used in MATLAB. Briefly, the underlying distribution for each time point was assumed to be log normally distributed. For each of the 1000 simulations in the bootstrap, linear regression was used to fit the sample to an exponential decay model, yielding a distribution of half-lives.*p* value was determined using a Student’s t-test on GraphPad Prism software.

### ELISpot Assay

The frequency of antigen-specific IFN-γ-producing CD8^+^ T cells was evaluated using the ImmunoSpot Mouse IFN-γ ELISpot Kit (Cellular Technology Limited). Briefly, spleen cells were harvested from immunized mice on Day 42, plated at 800,000 splenocytes/well and restimulated with 10 μg rabies virus glycoprotein peptide (AnaSpec) for 24h at 37 °C. Spots were developed following manufacturer’s instructions.

### Flow Cytometry Analysis of Immune Cell Infiltration

Briefly, hydrogels were excised from mice after euthanasia and were mechanically disrupted to create a cell suspension. For flow cytometry analysis, cells were blocked with anti-CD16/CD38 (clone: 2.4G2) and then stained with fluorochrome conjugated antibodies: CD3, CD11b, CD11c, CD19, CD45, CD335, Ly6G, Ly6C, and live-dead staining. Cells were then washed, fixed, and analyzed on an LSRII flow cytometer. Data were analyzed with FlowJo 10. See Table S1 for the antibody panel.

### Rabies Virus Neutralization Assay

Rabies virus neutralizing antibodies were measured using the Rapid Fluorescent Foci Inhibition Test (RFFIT). The test was performed by the Kansas State Veterinary Diagnostic Laboratory. Briefly, a series of serum dilutions are mixed with a standard amount of live rabies virus and incubated. Cultured cells are then added and incubated together with the virus and serum. Endpoint titer is calculated from the percent of virus infected cells at the end of incubation. Adequate protection was determined as rabies virus neutralizing antibody potency above 0.5 IU/mL, according to World Health Organization (WHO) guidelines.

### Animal Protocol

All animal studies were performed in accordance with National Institutes of Health guidelines and with the approval of the Stanford Administrative Panel on Laboratory Animal Care (APLAC-32109).

### Statistics

All results are expressed as mean ± standard error (SE) unless specified otherwise. Sample size for each experiment is included in the figure captions. Statistical analyses where more than two treatments were being compared were performed as general linear models in JMP Pro version 15. Post-hoc Tukey HSD tests for multiple comparisons were applied when treatment group was a significant fixed effect, and adjusted *p* values were reported. Statistical analyses where only two treatments were being compared were performed with a Student’s T test using Graph Pad Prism.

## Supporting Information

Supporting Information is available from the Wiley Online Library or from the author.

## Data Availability Statement

The data that support the findings of this study are available on request from the corresponding author. The data are not publicly available due to privacy or ethical restrictions.

## Acknowledgements

We would like to thank all members of the Appel lab for their useful discussion and advice throughout this project. Also, we are grateful to the staff of the BioE/ChemE Animal Facility who cared for our mice. We would also like to thank the Kansas State Veterinary Diagnostic Laboratory for their assistance with the RFFIT assay. We would like to thank the Stanford Shared FACS Facility for their assistance with data collection. Flow cytometry analysis for this project was done on instruments in the Stanford Shared FACS Facility obtained using NIH S10 Shared Instrument Grant (S10RR027431-01; 1S10OD026831-01). This research was financially supported by the Center for Human Systems Immunology with the Bill & Melinda Gates Foundation (OPP1113682; OPP1211043; INV027411). We thank the National Science Foundation Graduate Research Fellowship (JY and OMS), Eastman Kodak Fellowship (BSO), the Hancock Fellowship of the Stanford Graduate Fellowship in Science and Engineering (OMS), and the Stanford NIH T32 Biotechnology Training Grant (5T32GM008412-25; ELM).

## Author Contributions

JY and EAA designed the broad concepts and research studies. JY and EAA designed specific experiments. JY, BSO, ELM, OMS and NE performed research and experiments. JY and EAA wrote the paper. BSO edited the paper.

