## Supplemental Information for "A regimen compression strategy for commercial vaccines leveraging an injectable hydrogel depot technology for sustained vaccine exposure"

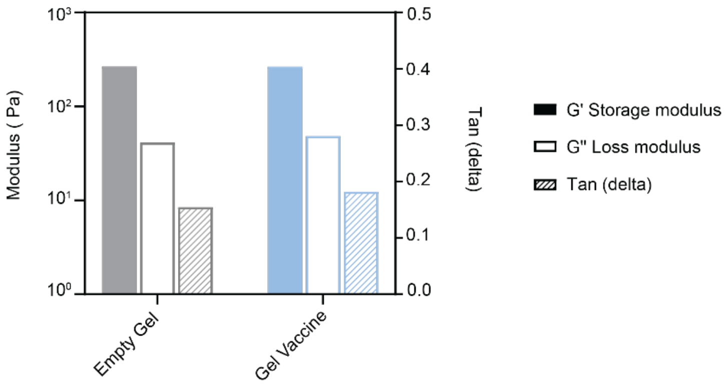


**Figure S1.** Mechanical properties of plain PNP-1-10 hydrogels and vaccine-loaded PNP-1-10 hydrogels at $\omega = 10$ rad/s.

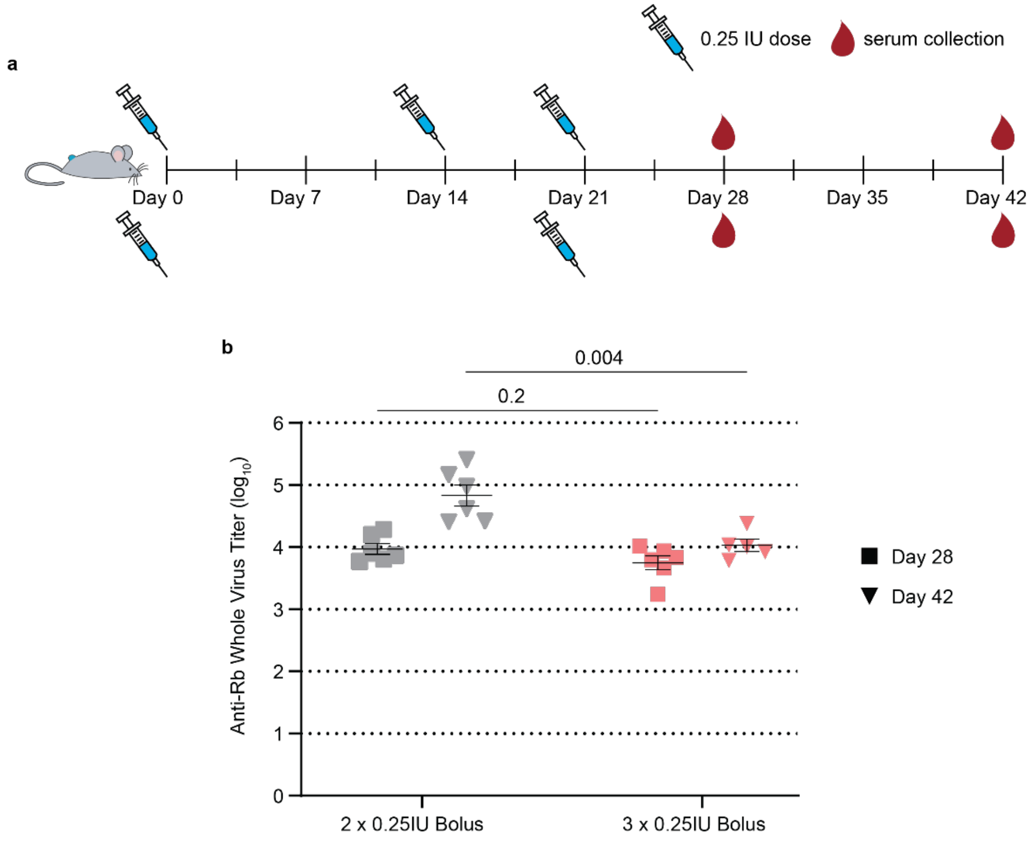


**Figure S2. Antibody titers following two and three dose immunization regimens of the bolus commercial RabAvert vaccine. (a)** Timeline of mouse immunizations and blood collection over a 6-week period. The two-dose regimen included immunizations on day 0 and day 21 while the three-dose regimen included immunizations on day 0, 7, and 21. **(b)** ELISA antibody titers for day 28 and day 42 after the entire immunization series. Data are shown as mean +/- SEM. *p* values were determined using a Student's t-test.

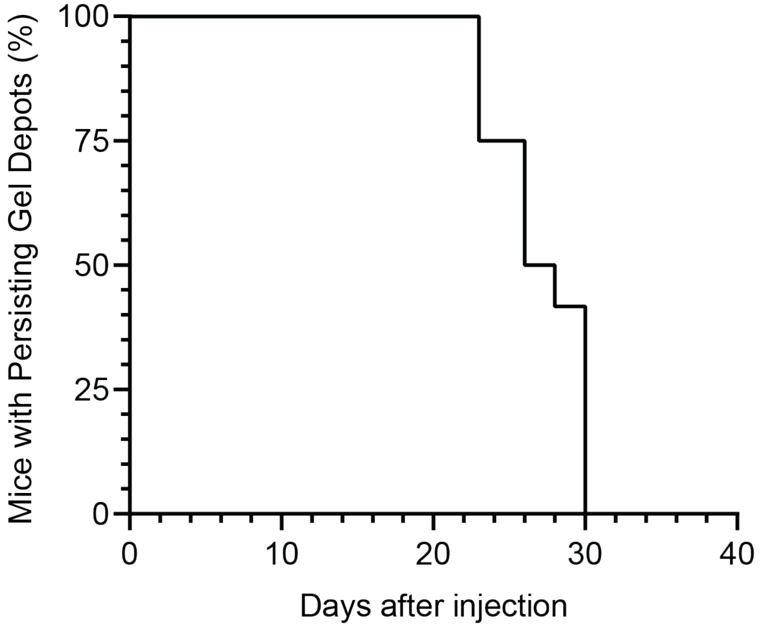


**Figure S3. Vaccine-loaded hydrogel depot persistence time in the subcutaneous space following administration by transcutaneous injection.** Mice were injected with 0.25 IU PNP-1-10 hydrogel-based vaccines (n = 12) and depots were tracked over time until no longer visible.

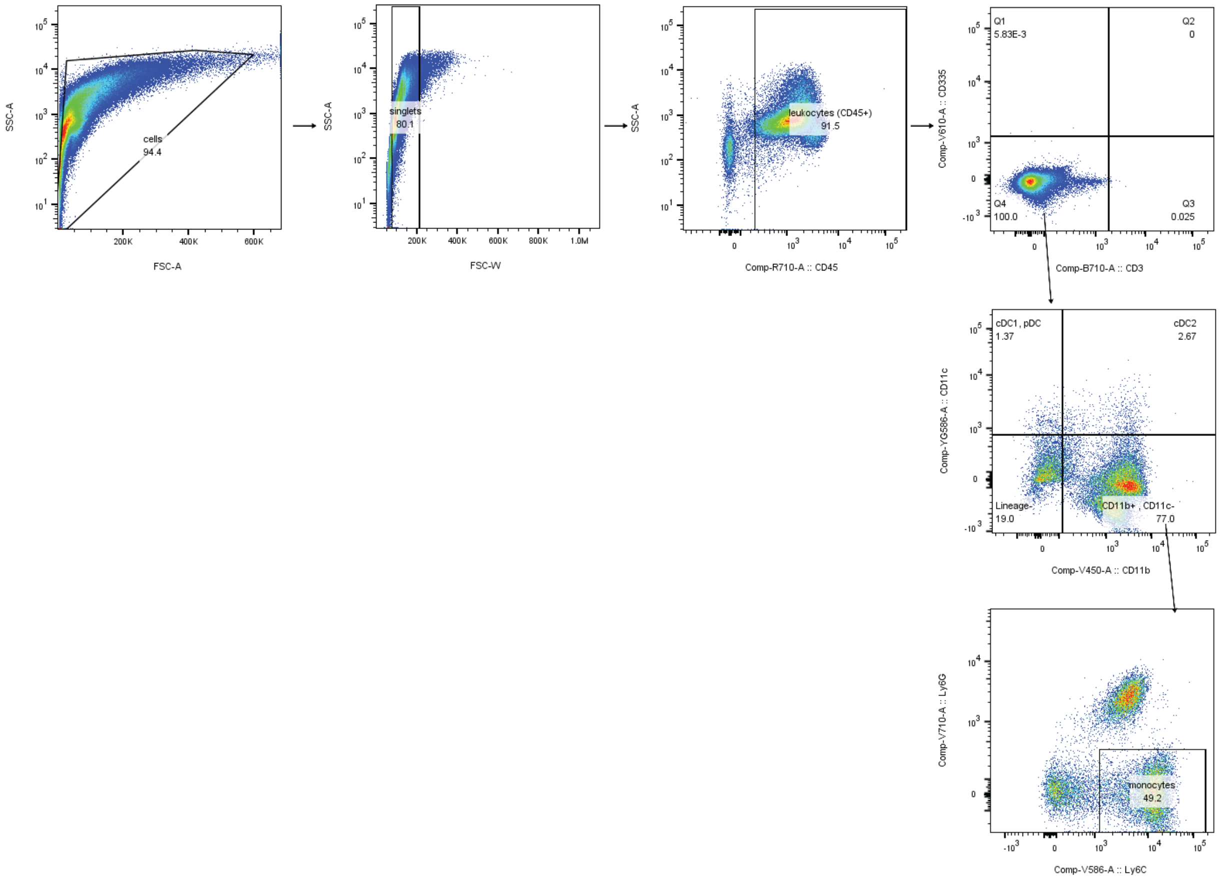


**Figure S4. Sample gating scheme for hydrogel niche.** Table S1 lists all fluorophores used for gating and analysis.

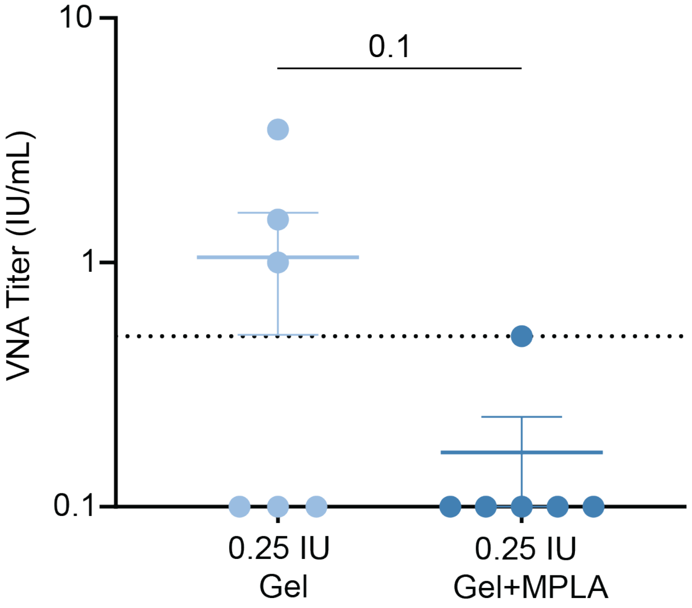


**Figure S5. Viral neutralizing antibody (VNA) titers.** At day 42 following immunization with PNP-1-10 hydrogel-based vaccines with and without MPLA adjuvant VNA titers were determined (n = 6). The dotted line represents the threshold of complete protection (0.5 IU/mL) as stated by the World Health Organization.

**Table S1.** Flow cytometry antibody information.

| Antibody (all anti-mouse) | Manufacturer | Clone |
|---|---|---|
| CD3 | Invitrogen | 500A2 |
| CD11b | BioLegend | M1/70 |
| CD11c | BioLegend | N418 |
| CD19 | BioLegend | 6D5 |
| CD45 | BioLegend | 30-F11 |
| CD335 | BioLegend | 29A1.4 |
| Ly6G | BD Biosciences (custom-made) | |
| Ly6C | BioLegend | HK1.4 |
| Live/Dead Near-IR | Invitrogen | Cat#L34975 |
